# Single-gene resolution of locally adaptive genetic variation in Mexican maize

**DOI:** 10.1101/706739

**Authors:** Daniel J Gates, Dan Runcie, Garrett M. Janzen, Alberto Romero Navarro, Martha Willcox, Kai Sonder, Samantha J. Snodgrass, Fausto Rodríguez-Zapata, Ruairidh J. H. Sawers, Rubén Rellán-Álvarez, Edward S. Buckler, Sarah Hearne, Matthew B. Hufford, Jeffrey Ross-Ibarra

**Affiliations:** Department of Evolution and Ecology and Center for Population Biology, University of California Davis, Davis, CA 95616; Department of Ecology, Evolution, and Organismal Biology, Iowa State University, Ames, Ia 50011; Institute for Genomic Diversity, Cornell University, Ithaca, NY 14853; Centro International de Mejoramiento de Maiz y Trigo (CIMMYT), Carretera Mexico-Veracruz, El Batan, 56130, Mexico; Department of Molecular and Structural Biochemistry, North Carolina State University, Raleigh, NC; Laboratorio Nacional de Genómica para la Biodiversidad/Unidad de Genómica Avanzada, Cinvestav, Irapuato, México; Program in Genetics, North Carolina State University, Raleigh, NC 27695-7614, USA; Department of Plant Science, The Pennsylvania State University, State College, PA, USA; Genome Center, University of California Davis, Davis, Ca 95616

## Abstract

Threats to crop production due to climate change are one of the greatest challenges facing plant breeders today. While considerable adaptive variation exists in traditional landraces, natural populations of crop wild relatives, and *ex situ* germplasm collections, separating adaptive alleles from linked deleterious variants that impact agronomic traits is challenging and has limited the utility of these diverse germplasm resources. Modern genome editing techniques such as CRISPR offer a potential solution by targeting specific alleles for transfer to new backgrounds, but such methods require a higher degree of precision than traditional mapping approaches can achieve. Here we present a high-resolution genome-wide association analysis to identify loci exhibiting adaptive patterns in a large panel of more than 4500 traditional maize landraces representing the breadth of genetic diversity of maize in Mexico. We evaluate associations between genotype and plant performance in 13 common gardens across a range of environments, identifying hundreds of candidate genes underlying genotype by environment interaction. We further identify genetic associations with environment across Mexico and show that such loci are associated with variation in yield and flowering time in our field trials and predict performance in independent drought trials. Our results indicate that the variation necessary to adapt crops to changing climate exists in traditional landraces that have been subject to ongoing environmental adaptation and can be identified by both phenotypic and environmental association.

## Introduction

Protecting crop and wild plant populations against the harmful effects of climate change is one of the most important goals of plant genetic research today. Temperature increases of 1.5°C above preindustrial levels are already occurring (1) with further increases of 4-7°C likely in the next 100 years (2). Agricultural systems have already been impacted by warming (3), and models under future warming scenarios indicate this trend will be exacerbated, with yield losses across major staple commodities in the range of 25-50% by 2100 (4).

One promising strategy for maintaining crop productivity under climate change is to harness existing genetic variation (5). Traditional domesticated varieties (*i.e*., landraces) and the wild relatives of many crops have been evolving in diverse environments for thousands of years (6; 7; 8), resulting in a broad base of genetic variation and adaptation to a wide range of environmental niches. Breeders have nonetheless been reluctant to use landrace and wild relative resources because incorporation of novel diversity requires many cycles of backcrossing and phenotyping to minimize linkage drag (9). But recent advances in breeding techniques such as rapid cycle genomic selection and molecular approaches including CRISPR/Cas9 now allow targeted transfer of alleles, saving both time and cost in breeding (10; 11) and present novel opportunities to take advantage of novel adaptive alleles. Transfer in this manner, however, requires detailed knowledge of candidate adaptive loci with precision to the gene or even nucleotide level. A significant impediment to this approach, then, is our limited understanding of the genetic basis of adaptation in most crops and their wild relatives.

Adaptive variation can be identified in crops by assembling germplasm sampled across gradients of interest, evaluating field trials across environmental extremes, and identifying population genetic signatures consistent with adaptation. Farmer seed networks and *ex situ* conservation efforts have preserved extensive collections of wild and landrace germplasm (12). Adaptation in such materials can be identified through evaluation in observation trials (common gardens) and reciprocal transplant experiments that evaluate variation in fitness components across environments. Maize landraces grown across in common gardens across elevational gradients in Mexico, for instance, demonstrated substantially higher fitness when grown closer to their natal environments (13). Population genetic signatures of adaptation can also be identified using correlations between environmental gradients and allele frequencies (14; 15; 16; 17). For example, (15) showed that Sorghum alleles associated with drought and low pH can be used to predict plant phenotype, and (18) predicted *Medicago* growth in controlled environments using environmentally associated SNPs. While there is great promise for identifying adaptive variation for crop improvement using these combined approaches, comprehensive efforts are lacking.

Maize is the most productive crop globally and has adapted to grow in nearly every environment that humans inhabit (19), but yield in modern maize lines has already been impacted by climate change (20). Maize landraces and its wild relatives the teosintes are well adapted to their distinct local environments (6; 21) and constitute an extensive but largely untapped well of genetic diversity (22; 23). CIMMYT (the international maize and wheat improvement center), which is part of the CGIAR network, curates the world’s largest collection of maize landrace accessions (>24,000), a resource that could play a crucial breeding role in mitigating the negative effects of climate change on maize.

In this study, we identify adaptive genetic variation in a core set of CIMMYT’s global collection of maize landraces based on the results of field trials replicated across a range of environments and a population genetic analysis that identifies loci associated with environmental variation across the landscape. We demonstrate that alleles exhibiting genotype by environment (*G* × *E*) interactions for fitness across trials are predictably associated with climatic variables spanning the range of landraces in Mexico. Likewise, we show that environmental association (eGWAS) can identify candidate genetic loci which are linked to phenotypic and fitness differences across trials. We show substantial overlap in the loci identified by these analyses; alleles at loci associated with fitness difference between trials tend to be found in their beneficial environments. We validate a set of loci associated with precipitation by demonstrating their effect on fitness in independent drought trials, and identify a list of candidate genes from both approaches that also show population genetic evidence of recent selection.

## Results

To identify patterns of local adaptation in maize landraces we assayed plant phenotypes across 23 trials in 13 different common garden field locations in Mexico (Fig. 1A). Briefly, each of more than 2,200 landrace individuals was crossed to a locally adapted hybrid tester and the F1 progeny were grown at each site, resulting in a total of approximately 280,000 plants with phenotypic data (see Methods). Maize plants in these common gardens exhibit a classic phenotypic pattern of local adaptation: plants are taller, flower earlier, and exhibit higher yield (measured as field weight, bare cob weight, or grain weight per hectare) and less stress (lower anthesis-silking interval) when grown at elevations closer to their original collection locality (Fig. 1B; *P* < 1e^-06^ for each phenotype). We observe similar patterns for other environmental variables including mean temperature, precipitation, isothermality, and the number of wet days an experimental trial receives (Supplemental Figs. S4-S7), suggesting broad adaptation to local environments, consistent with previous transplant experiments (24; 25).

**Figure 1.**
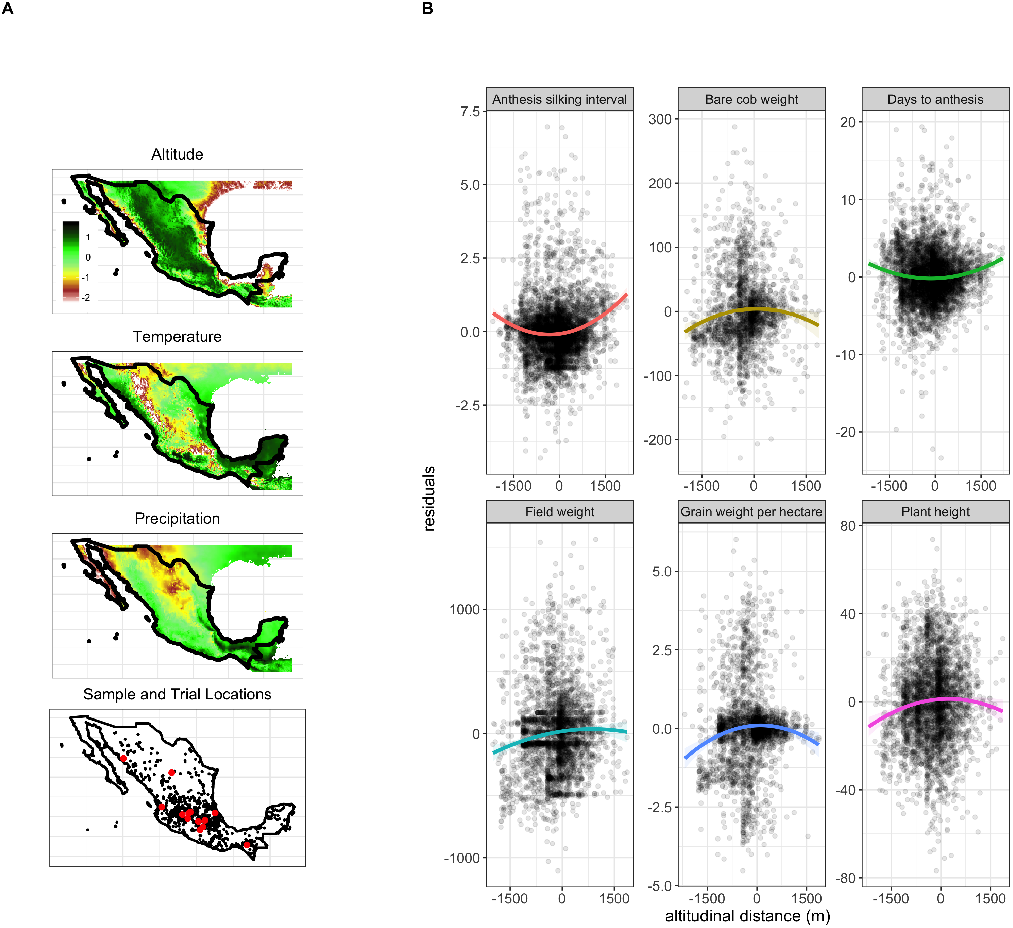
Maize plants grown in experimental gardens are adapted to diverse environments. A) Temperature, elevation, and precipitation across Mexico (Z-standardized). The bottom plot shows locations that landraces originated from (black) and the location of phenotyping trials (red). B) Fitness decreases as elevational distance between common garden and collection locality increases.

To identify loci underpinning this strong genotype by environment (*G* × *E*) effect on plant fitness, we implemented a mixed linear model in GridLMM (26) to perform an association analysis of fitness-related phenotypes (field weight, grain weight per hectare, bare cob weight) across our study sites (Fig. 2A; see Methods). For each of the top 1000 SNPs, showing significant *G* × *E*, we polarize the major allele by the field trial environment in which it increases fitness. This polarization accurately predicts where alleles are naturally found across the landscape: alleles that increase fitness in high elevation trials, for example, are found roughly 100-400 meters higher than random set of frequency-matched alleles and 400-600 meters higher than alleles that increase fitness at low elevations (Fig. 2B). In addition to predicting the modern geography of maize variants, our top *G* × *E* loci also predict allele frequency differences between distinct locally adapted teosinte subspecies. Lowland (*Zea mays* ssp. *parviglumis*) and highland (*Zea mays* ssp. *mexicana*) teosinte diverged 60,000 years ago (27) and subsequently adaptated to distinct ecological niches (28). Yet we show that long-term ecological divergence between these wild subspecies is predictable from the *G* × *E* effects of an allele in maize (Fig. 2C), suggesting either selection has maintained such alleles as standing variation over relatively long periods or gene flow has transferred adaptive alleles into maize (29).

**Figure 2.**
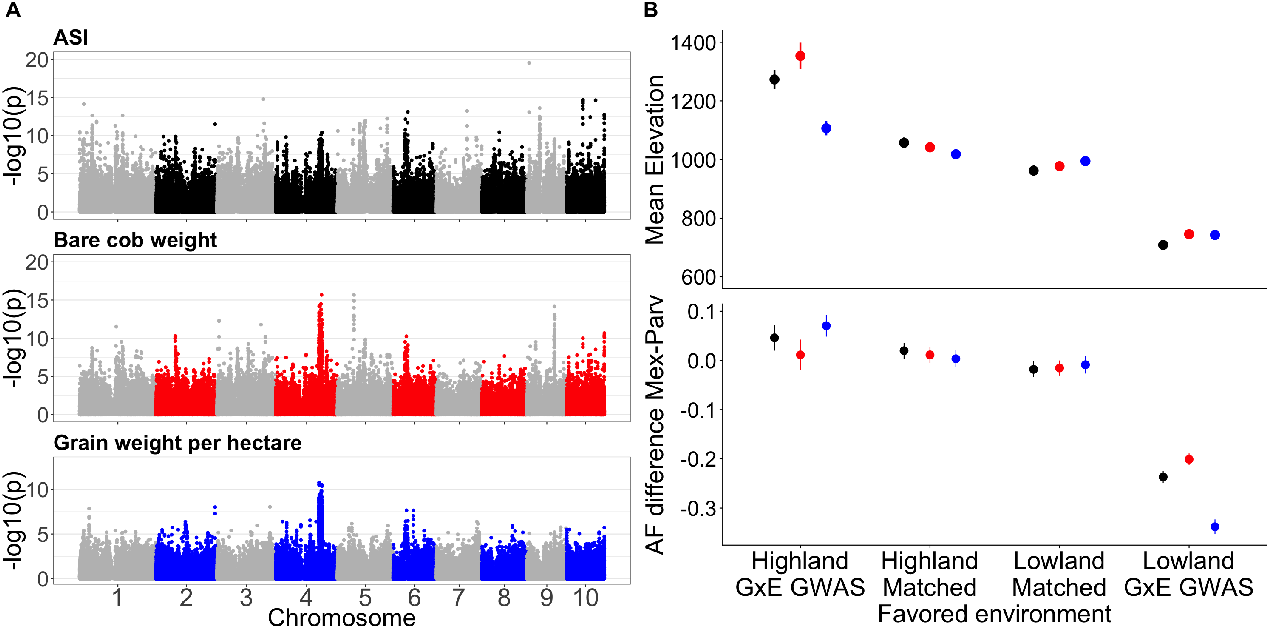
Genotype by environment effects predict SNP association with the environment and frequency in wild populations. A) Genome-wide association of G × E effects for three fitness-related traits across all common gardens. B) Alleles that are favored in high elevation trials exist at higher elevations across Mexico (top) and show a larger frequency difference between the highland teosinte *Zea mays* ssp. *mexicana* and its lowland counterpart *Zea mays* ssp. *parviglumis* (bottom). Colors represent the phenotypes in part A. Lines extending above and below points represent 95% confidence intervals. In both plots, SNPs showing significant GWAS effects are compared to a control set with matched allele frequencies.

One of the clearest features of our *G* × *E* association is a broad peak on the long arm of chromosome 4 that is identified for several distinct phenotypes (Figs. 2A and S8). This region harbors the well-characterized inversion polymorphism *Inv4m* which spans ≈ 15Mb. The non-reference haplotype at *Inv4m* appears to have arisen via introgression from the highland teosinte subspecies *Z. mays* ssp. *mexicana* (29), is associated with a number of adaptive phenotypes (29; 30), and shows strong population genetic signals of selection (31). Although large-scale polymorphisms like *Inv4m* often stand out, a key advantage of our association approach is its high resolution — 41% contain a single gene, and nearly 80% contain fewer than three genes. A detailed description of association peaks and potential candidate genes can be found in Supplemental Tables S1-S6.

Variation in lipid composition has been linked with multiple environmental factors (32; 33; 34; 35; 36; 37), is associated with variation in flowering time (38), and population genetic analysis suggests that lipid metabolic pathways may have been important in local adaptation to highland regions in Mexico and South America (31). We thus compared our results to a QTL analysis of *G* × *E* for phospholipid metabolites in Mexican highland maize (39). We find a number of specific lipid *G* × *E* QTLs that overlap with loci we identify as showing *G* × *E* flowering time and yield across elevation (Supplemental Table S7). Among the overlapping regions, we found three genes (Zm00001d043228, Zm00001d043207 and Zm00001d039542) involved in the synthesis and degradation of phospholipids. All three are up-regulated after cold stress in temperate inbreds (40) (Supplemental Fig. S9).

While our *G* × *E* associations directly test the link between fitness and genotype across many trial environments, environmental association offers even greater climatic precision. Using genotype data from nearly 4,000 geo-referenced maize landraces across the Americas (Fig. 1A 30), we evaluated the association between SNP genotype and both 50-year average climate (41) and fine-scale measurements from local weather stations over the course of the local growing season (see Methods). As with our previous phenotypic *G* × *E* results, our environmental associations (eGWAS) highlight *Inv4m* but nonetheless exhibit a similar fine-scale resolution (41% of association peaks contain a single gene, while only 20% contain more than two genes).

Loci identified in our environmental association (Supplemental Figs. S10, S11 show strong *G* × *E* effects on yield, increasing plant survival and reproduction in trials most similar to their associated environment (Fig. 3B); this effect also holds for a number of relevant phenotypes (Fig. 3C). But while eGWAS SNPs show strong *G* × *E* for fitness, we see little difference in their average main effect S13, as expected if such alleles are favorable only in a limited number of environments. In contrast, we see only a very weak *G* × *E* effect of eGWAS SNPs on flowering time (Fig. 3D), but a large main effect. While selection will quickly fix or remove variants with large average effects on fitness, variants with large effects on flowering time or other phenotypes can segregate at appreciable frequency (30) if the phenotypic optimum varies across the species’ range (e.g. 42; 43).

**Figure 3.**
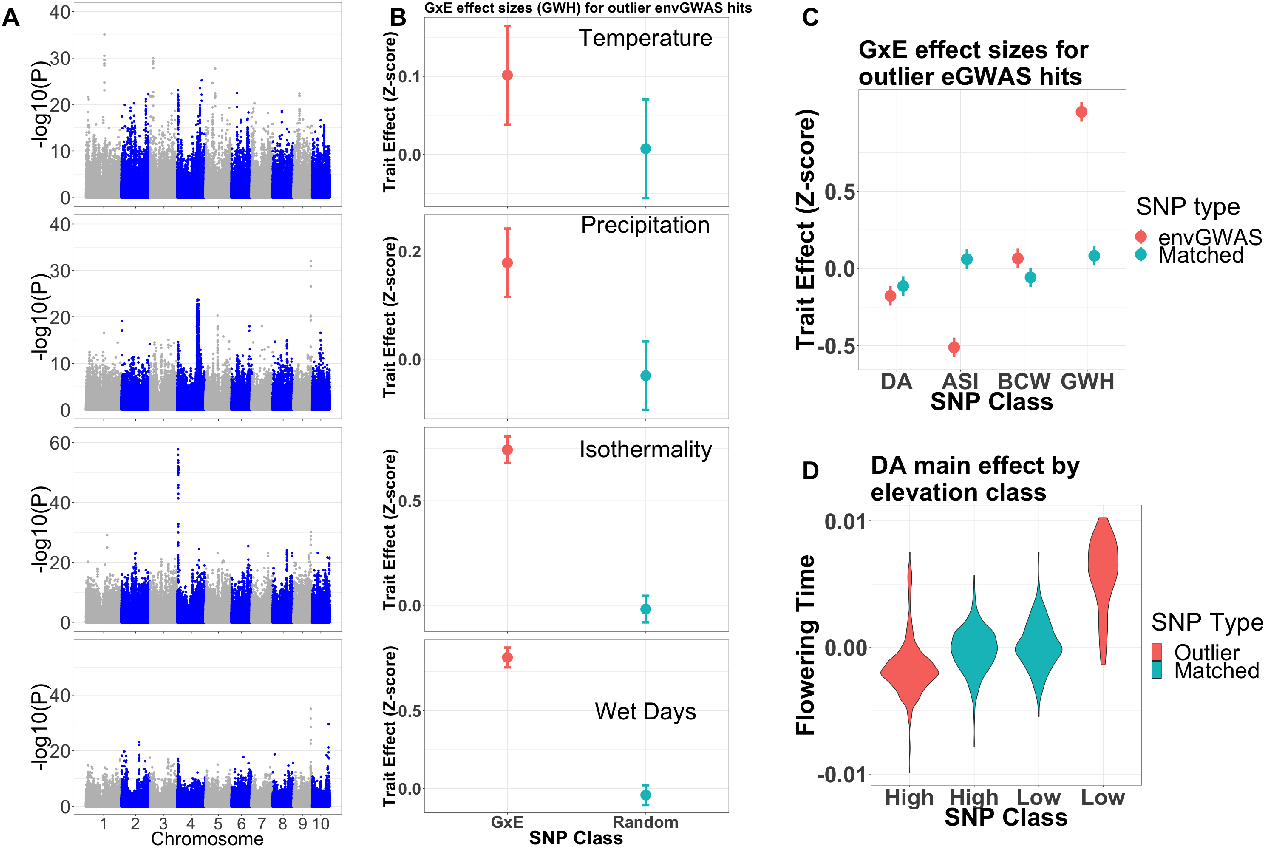
Adaptive variation from environmental GWAS. A) Manhattan plots showing associations with temperature, precipitation, isothermality, and the number of wet days in the growing season. B) SNPs identified via environmental association show strong *G* × *E* effects on grain weight per hectare, increasing fitness in common gardens more similar to their environment of origin. C) Environmental GWAS hits across elevation predict beneficial *G* × *E* of DA: days to anthesis, ASI: anthesis-silking interval, BCW: bare cob weight, and GWH: grain weight per hectare (top). D) Additional levels of adaptive flowering time are influenced by main effect as environmental GWAS alleles originating from lowlands flower later than their highland counterparts (and vice versa).

To validate our eGWAS analyses, we tested whether alleles associated with annual precipitation would impact yield in a pair of independent drought trials including more than 1,000 of our genotyped accessions (see Methods). We find that alleles associated with low precipitation across the landscape are more likely to increase yield in drought conditions than either high precipitation alleles or alleles from randomly sampled frequency-matched loci (*P* < 1e^-7^, Supplemental Figure S3. We combined information from this drought experiment with expression data of leaf and ovary response to drought (44) in order to further refine candidate drought adaptation genes (Figure 4; a full list is in Supplemental Tables S8-S10). For example, 6 SNPs in Zm00001d003176 (Nodule inception protein-like protein 3; *nlp3*) are associated with precipitation across Mexico and 3 are associated with yield in our drought trials. Interrogation of the expression data shows that *nlp3* also shows significant gene expression differences between well watered and drought conditions (*P* = 7.3e^-05^). *nlp3* acts as a central signaling protein to control nitrate and phosphate responsive genes (45), is responsive to multiple environmental stress conditions, and *nlp3* responses change carbon, nitrogen, and phosphorous metabolism (46). Another candidate is Zm00001d002999 (textitga2ox2), which functions in giberellin catabolism. Consistent with our expression analysis (Figure 4), textitga2ox2 is upregulated in areas of active growth during drought stress (47), leading to a reduction in gibberellic acid that slows elongation and in turn allows for resources to be allocated to other drought dependent processes (48). Zm00001d004621 (*ntr1*) is a peptide transporter and another clear for drought adaptation — when overexpressed in soybean it increased seedling adaptation to drought (49). Finally, candidate gene Zm00001d046621 is *glossy15*, an apetala2 type transcription factor. Variation at textitglossy15 hastens or delays the transition from juvenile to adult stage (50; 51; 52), which will influence drought avoidance ability as the transition from juvenile to adult stage is required for flowering in maize. Additionally, *glossy15* variation influences the composition and amount of epicuticular waxes on young maize leaves (53) which can reduce drought tolerance in maize seedlings (54). Finally, we note that drought expression data further strengthens candidates identified previously — all three of the candidates highlighted as showing *G* × *E* for both phospholipid content and phenotypes additionally show differential expression under drought as well as cold conditions (Supplemental Fig. S9).

**Figure 4.**
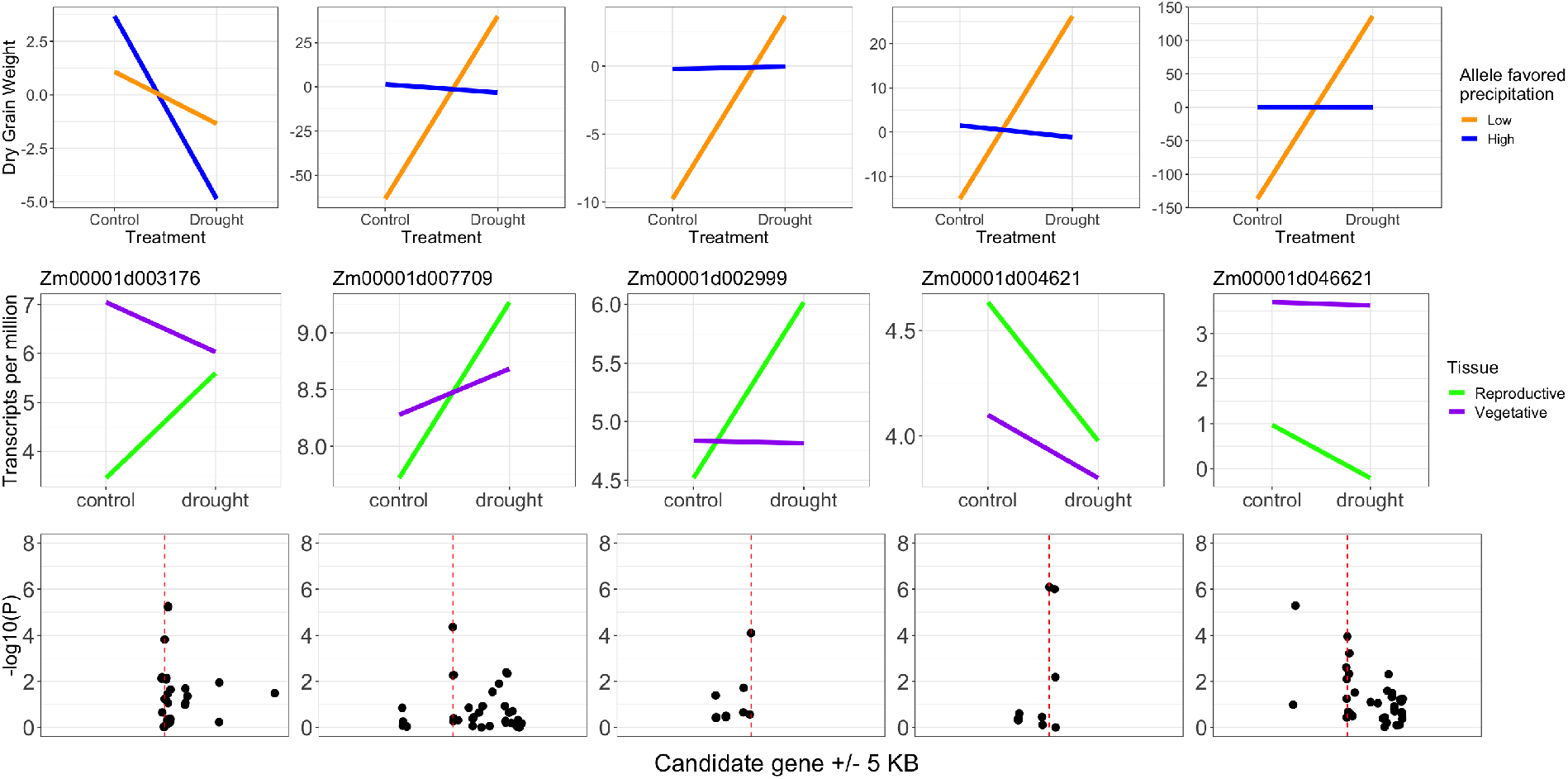
Environmental GWAS identifies candidate drought improvement genes. Candidate SNPs (columns) have low precipitation associated alleles that increase yield in drought trials (top row) and show differential expression under drought (middle row). Significant environmental GWAS (precipitation) p-values for each gene are highly localized and do not extend to nearby genes. Manhattan plots (bottom row) show the precision of environmental GWAS around each candidate SNP (broken red line) and 5KB upstream and downstream of each candidate. Each candidate has no other gene models present within the extent of the windows.

We also compared out results to published yield data from drought and heat stress trials in the “Drought Tolerant Maize of Africa” (DTMA) panel (55). We compared our top 5000 eGWAS hits to *G* × *E* associations with yield across environments in the DTMA, finding that 142 of our eGWAS hits are associated with significant (*P* < 1e^-3^) drought-specific yield effects in one of the three measured traits (Supplemental Tables S11-S13). At the end of chromosome 9, for example, is the strongest eGWAS hit for precipitation and wet days during the growing season (Figure 3A) as well as diurnal temperature range and isothermality S11. Although this association peak includes roughly 7 different genes, an annotated heat-shock-related transcription factor (hsftf9; Zm00001d048041) is a compelling candidate, as its *Arabidopsis* ortholog (AtHSF1) is expressed under heat stress and is responsible for activation of heat-shock-protein-based thermotolerance (56; 57), while its tomato homolog HSFA1 acts as a master regulator of thermotolerance (58). Indeed, a SNP in the 5’ UTR of hsftf9 showed a strong G × E association for yield between drought and well watered treatments (p=0.00012) in the DTMA drought panel. Another candidate gene identified from our eGWAS (Zm00001d007709; Figure 4) is from a family of proteins containing In Between Ring (IBR) domains, and likely functions in protein degradation through protein-protein interaction. Little is known about the function of this gene in maize or of its most similar relative (ATARI7) in *Arabidopsis*. But despite the lack of functional information around this gene, the SNPs tested within the coding sequence of this gene show at least one significant (*P* < 0.01) hit for all three traits (DA, ASI, and Yield) tested in the DTMA drought trials.

To highlight a combined set of candidates for characterization, we complemented our *G* × *E* and environmental associations with a phenotype- and environment-independent scan for SNPs showing extreme patterns of allele frequency differentiation across the landscape (see Methods). We focused our assessment of gene function on the set of 39 genes (Supplemental Table S14) identified as candidates in all three approaches (*G* × *E*, eGWAS, and PCAdapt), many of which have functions that consistent with adaption to abiotic and biotic stress. Zm00001d046909 (homogentisate phytyltransferase *hpt1*), for example, is active in Vitamin E biosynthesis (59; 60), known to play a role in tolerance to salinity, drought, cold temperature, metal toxicity, ozone, and UV radiation stress (reviewed in (61)). SNPs in Zm00001d046909 are associated with daily temperature variability during the growing season, and show *G* × *E* effects for ear mass and temperature, precipitation, and elevation. The candidates Zm00001d024909 (CCT domain transcription factor) and Zm00001d010752 (*pebp8, zcn8*) are known to be involved in photoperiod sensitivity, flowering time, and long-day stress response (62; 63; 64). The former shows environmental association with mean temperature and is a candidate in our *G* × *E* analysis between ear mass and temperature, precipitation, and elevation, and the latter is a *G* × *E* candidate for flowering time and temperature as well as annual precipitation and wet days). Finally, we identify Zm00001d045972 (*umc81*) as a candidate for *G* × *E* between yield and elevation, and as showing allele frequency associations with temperature and water vapor pressure, but it has been previously highlighted as a candidate for days to anthesis (65), plant height (66), drought stress tolerance (67), and root hair length plasticity (68).

## Conclusions

Developing diverse crops that can deliver increased yields in the face of a changing global climate is one of the greatest challenges facing modern agricultural research. Understanding and characterizing adaptive loci will allow for the incorporation of novel variation into breeding efforts to adapt crops to changing climates. Although *in situ* breeding remains an important source of adaptive variation (6), our work highlights the utility of leveraging *ex situ* germplasm collections to evaluate phenotype across a diverse set of environments. Such collections typically have location information which can further increase the value of genomic data by information about the environments to which taxa have been locally adapted (69; 18; 15; 70; 14; 16; 30; 17). We use these data to identify loci underlying phenotypic and fitness differences in maize across environments and demonstrate that our environment-specific effect estimates successfully predict where alleles are found across the landscape. We then show that approaching the problem from the other direction — identifying loci that associate with environmental variables — can precisely delineate individual genes that show predictable phenotypic effects in field trials. In sum, we have validated the utility of phenotypic and environmental association approaches for identifying relevant adaptive variation and provided a high-resolution list of hundreds of candidate genes showing environmental and phenotypic associations that can be directly tested in targeted experiments (e.g. CRISPR-mediated knockout 71; 72) to improve climate adaptation in modern breeding populations.

## Methods

### Samples and Genotyping

The Seeds of Discovery project (SeeD) has genotypically characterized the CIMMYT International Germplasm Bank maize collection containing over 24,000 landrace accessions primarily from North, Central, and South America. Here, we used data generated across a core collection of ≈ 4, 500 individual accessions from this project representing the breadth of environmental variation across the Americas as described in (30). Briefly, each individual was sequenced using genotype-by-sequencing (73) to a median 2X coverage. Genotypes were called using TASSEL (74), resulting in an initial set of 955,120 SNPs reported in (30). In addition to these data, we used GBS genotypes from the wild teosintes *Zea mays* ssp. *parviglumis* and *Zea mays* ssp. *mexicana* from (75). All genotype data were uplifted to coordinates on the B73 v4 reference genome (76) using the crossmap liftover tool (77).

### Multi-Environment Estimation of Phenotypes

As part of the SeeD evaluation of the landrace core collection, offspring of each of the genotyped individuals were planted in multiple environments under a replicated F1 crossing design (see 30, for design details). Two important features of the crossing experiment were added to ensure that phenotype data was not overly biased by elevational adaptation. First, plants were preferentially grown in locations that were of similar adaptation (highland tropical, sub-tropical or lowland tropical) to their home environment. While F1s with a highland accession parent were grown in low elevation and vice versa, on average, more plants from highland accessions were grown in high elevation than low elevation trials (Figure S2). Second, each plant was crossed to a test plant that was adapted to the environment that the F1 seeds were grown in. Both of these design features facilitates comparison of a larger sample but reduces adaptive differences among landraces and make our estimates of adaptive differences more conservative. The resulting F1 offspring were then grown in replicate in ambient field conditions across a total of 23 trials in 13 locations in 2 years. Trials used in the study are detailed in Supplemental Table S15, data is available from http://hdl.handle.net/11529/10548233.

Although the collections are from across the Americas, the overwhelming majority were sampled from the center of maize diversity in Mexico. Because these samples had the best climate data, and to minimize complications arising from large differences in plant environment of origin, trial environments, and population structure (31), we restricted our analysis of field phenotype data to the 2433 landrace accessions of Mexican origin. A range of 9-26 (average of 16) offspring per landrace cross were measured in an augmented row-column design to provide additional control for random stochastic variation; Figure S2 shows the number of accessions with phenotypes across the elevational range of the trials.

Across the 13 sites plants were phenotyped for a variety of traits. We used data from plant height (PlH), field weight (FW), bare cob weight (BCW), and moisture adjusted grain weight per hectare (GWH). In addition to these phenotypes, we added previously published data on flowering traits (days to anthesis (DA), days to silking (DS), and anthesis-silking interval (ASI)) from these same trials (30).

### Phenotypic Analysis

Breeding values for each landrace were estimated as described in Navarro *et. al*. (30), controlling for tester, checks, and field position in a complete nested model. We then removed any plots that had a reported disease score and also removed trials and accessions with outlying phenotype scores (typically > 4 standard deviations from the mean). Overall, there were 23 distinct trials between the 13 sites over 2 years, each with breeding values calculated for an average of approximately 850 accessions. The GxE phenotype datasets that we used for calculating GxE outliers ranged from 11,762 to 4,851 measurements depending on what trait was measured 1.

To test for associations between markers and responses to environmental variation across the trials (i.e. GxE effects) for each phenotype, we used the following mixed-effects GWAS model for each marker *i* = 1…*p*:

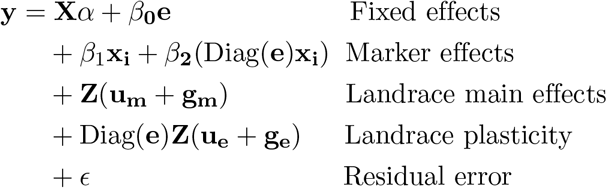

Here, **y** is a vector of *n* phenotypes (breeding values) for each landrace across all trials that it was tested in; **X** is a *n* × 8 design matrix specifying fixed effects for year (2) and testers (7) with coefficient vector *α*; **e** is a vector representing the environmental value, i.e. the elevation, mean temperature or precipitation of the trial, with mean effect *β*_0_; **x_i_** is a vector or genotypes at marker *i* for each observation and *β*_1_ and *β*_2_ are marker effects on the intercept and response to the environment, respectively; **Z** is a *n* × *r* design matrix for landraces, and **u_m_, g_m_, u_e_**, and **u_e_** are random effects accounting for the expected correlations of intercepts (*m*) and responses to the environment (*e*) among observations due to shared background additive (*u*) and non-additive (*g*) genetic variation for the multiple observations of the same landrace across different trials; and **e** is a vector of the environmental variable (i.e. mean phenotype value of the trial). The random effects have distributions:

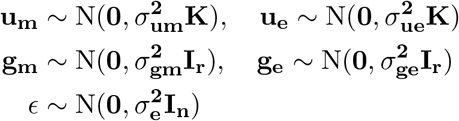

where **K** is the *r* × *r* kinship matrix for the *r* ≈ 2200 landraces computed in Tassel 5 (74) using a set of ≈ 6,500 unimputed loci that contain over 95% complete genotype data, **I_r_** is the *r* × *r* identity matrix and **I_n_** is the *n* × *n* identity matrix.

We used the `GridLMM_GWAS` function (26) to perform a GWAS scan across all markers, re-estimating the four variance components by REML separately at each marker (`method = “REML”`) using GridLMM-fast heuristic (`algorithm = “Fast”`) with `h2_step = 0.1`, and tested the significance of the marker main effects *β*_1_ and plasticity effects *β*_2_ independent by Wald tests.

### Environmental associations

In order to identify loci important for environmental adaptation, we performed an additional association analysis between allele frequency and environmental variables across the landscape. Here, to take advantages of a wider array of environmental variation, we expanded our sampling of individuals to all 4,127 genotyped accessions with latitude and longitude coordinates, including samples collected outside of Mexico. For general environmental layers (such as elevation, mean annual temperature, diurnal temperature range, mean isothermality, mean annual precipitation, and precipitation seasonality) we used latitude and longitude coordinates to infer values using the WorldClim 2 (41) layers at 0.5 minute resolution (https://data.cimmyt.org/dataset.xhtml?persistentId=hdl:11529/10548079) based on longterm averages for 1970-2000 for the climatic parameters as well as elevation data at 90 m resolution from the global elevation model from NASA and CIAT (78). In addition to these broader climate measurements, we included detailed weather station measurement based gridded datasets of the frequency of wet and frost-free days, cloud cover, vapor pressure covering the period 1950-2012 (79) and a longterm aridity index (80), each averaged over the approximate six month local growing season of accessions. Additional data sets representing soil pH (81), and soil waterlogging percentage (82; 83) were used to extract values to represent soil-based abiotic stresses in the environments where accessions originated. Due to differences in collection specificity, some individuals in the panel have unverified latitude longitude coordinates, and we restrict our detailed climate analyses to a curated subset of individuals (roughly 2,900 individuals) with verified location data.

For each different environmental layer we fit a model to measure the association between SNP genotype **x_i_** and the given environmental values for each accession **p**, correcting for population structure using the first five principal components **Q**:

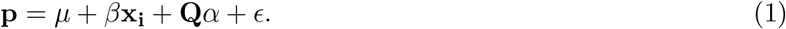

We then tested the significance of *β* with an F-test. In this analysis we only use loci with a minor allele frequency greater than 10%.

To determine if environmentally associated loci are contributing to adaption we test whether outlier loci contribute more to adaptive phenotypic differences in our experimental trials. We first selected 1,000 SNPs with the strongest association (lowest p-value) with elevation and a set of randomly selected SNPs that match the frequency deciles of associated SNPs. For each SNP in both sets, we used our field evaluation to estimate the standardized genetic value (in z-scores) of the main effect and the *G* × *E* term and tested if environmentally associated loci had a larger phenotypic effect than randomly matched loci. To ameliorate issue with sampling nonindependent SNPs, we excluded SNPs inside the large inversion *Inv4m* from this analysis.

### Gene Identification

To identify genes in our GWAS signal, we first smoothed the per-SNP association p-values across each chromosome using a spline analysis implemented in the R package GenWin (84). We used the maize reference B73 v4 annotation (76) to define gene positions and used the ‘countoverlaps’ function from the GenomicRanges R package (85) to determine the number and identity of genes that overlap with outlier SNP windows.

### Drought Trials

We also use drought trials to validate adaptive drought tolerance loci. The first drought experiment dataset we use was generated as part of the SeeDs of discovery phenotyping data discussed earlier and consists of 1,057 different accessions grown in drought and well watered trials in the Iguala and Obregon localities. These accessions were phenotyped for dry grain weight in drought and well-watered conditions at both locations, for a total of 2,047 yield phenotypes. We used GridLMM to fit a model with tester and location as covariates and measured the strength of the *G* × *E* yield response as above. In order to determine the effect of environmental GWAS loci on phenotypes in drought we ran an association analysis on the top 1,000 environmental association hits from annual precipitation and 1,000 randomly chosen loci matched for allele frequency. As above, we tested whether alleles associated with low precipitation would increase yield in drought trials in comparison to the alternate high precipitation alleles by making a prediction of the low and high precipitation genotypes from the model for both the environmentally associated and randomly paired loci.

### Drought Tolerant Maize in Africa

Three hundred maize lines from CIMMYT IITA (the International Institute of Tropical Agriculture) were previously genotyped and tested for drought tolerance in yield trials of well watered, drought, high heat, or combined conditions in the Drought Tolerant Maize for Africa (DTMA) GWAS panel (55). We used GBS genotypes for the DTMA from (86), allowing us to test associations between individual genes and phenotypes. We used GridLMM (26) to fit a *G* × *E* model analogous to that used previously in our phenotypic trials. In this model we fit grain weight per hectare of the as explained by genotype, the treatment the phenotype was collected in, and the interaction term (the change in yield across different trials).

### Expression analyses

To test for regulatory variation in candidate drought genes we downloaded raw RNA-seq data from (44). This experiment consists of replicated individuals of the inbred line B73 grown in drought and well watered conditions, with expression measured in the meristem of the three youngest leaves and ovary tissue at 1 day after pollination. We used kallisto (87) to estimate expression based upon kmer abundance of RNA-seq data at the gene models from the maize v4 genome. We used sleuth (88) to normalize and fit a factorial model to test the strength of the effect of growth condition and the growth condition by tissue interaction effect compared to a null model with only the main effect of tissue specific differences.

## Supplemental Tables and Figures

**Figure S1.**
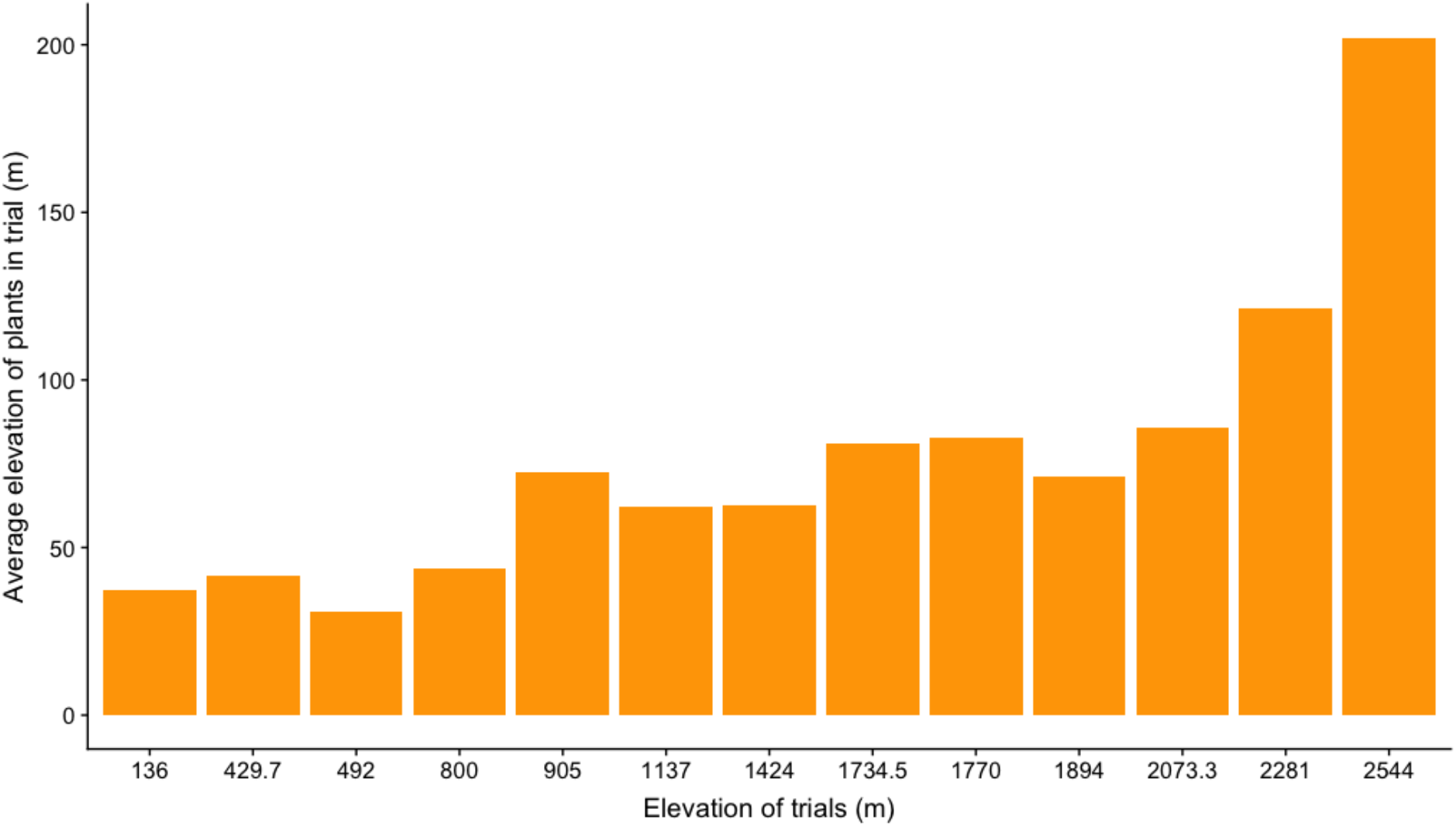
Average elevation of origin of plants grown in elevations at different trials.

**Table 1.**
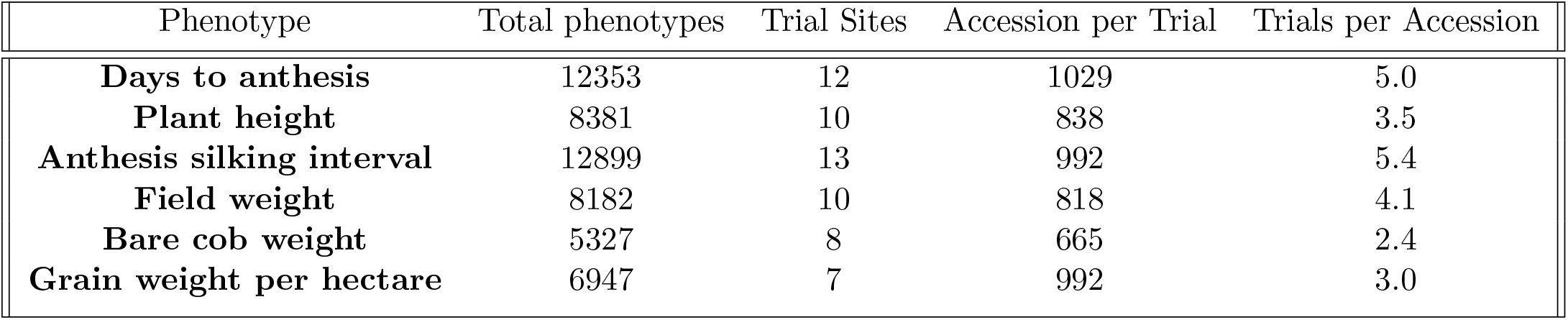
Number of accessions and trials of different traits

**Figure S2.**
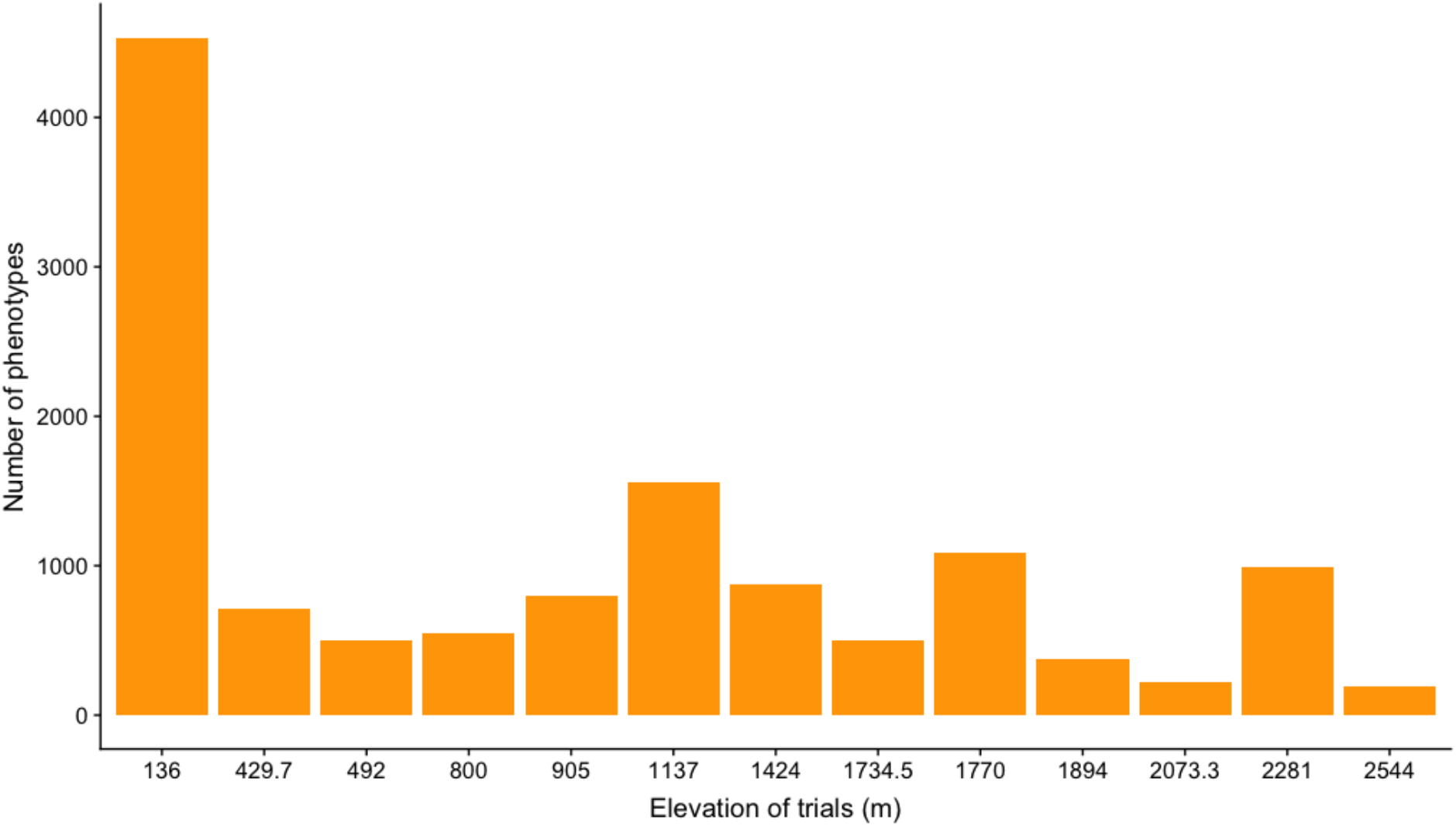
Number of phenotypes collected at different elevational classes

**Figure S3.**
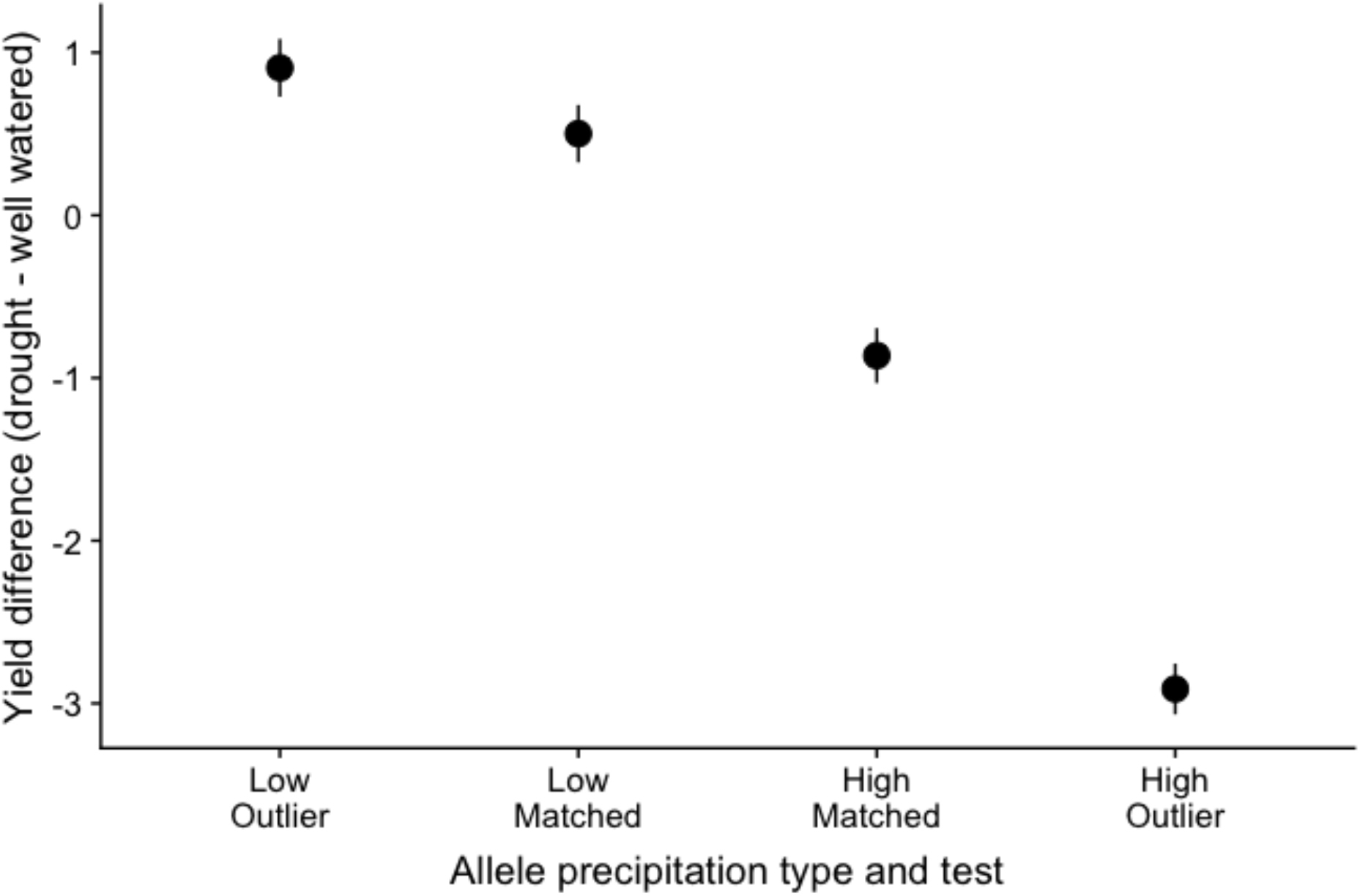
Yield performance in drought trials of precipitation outlier alleles categorized by environmental preference

**Figure S4.**
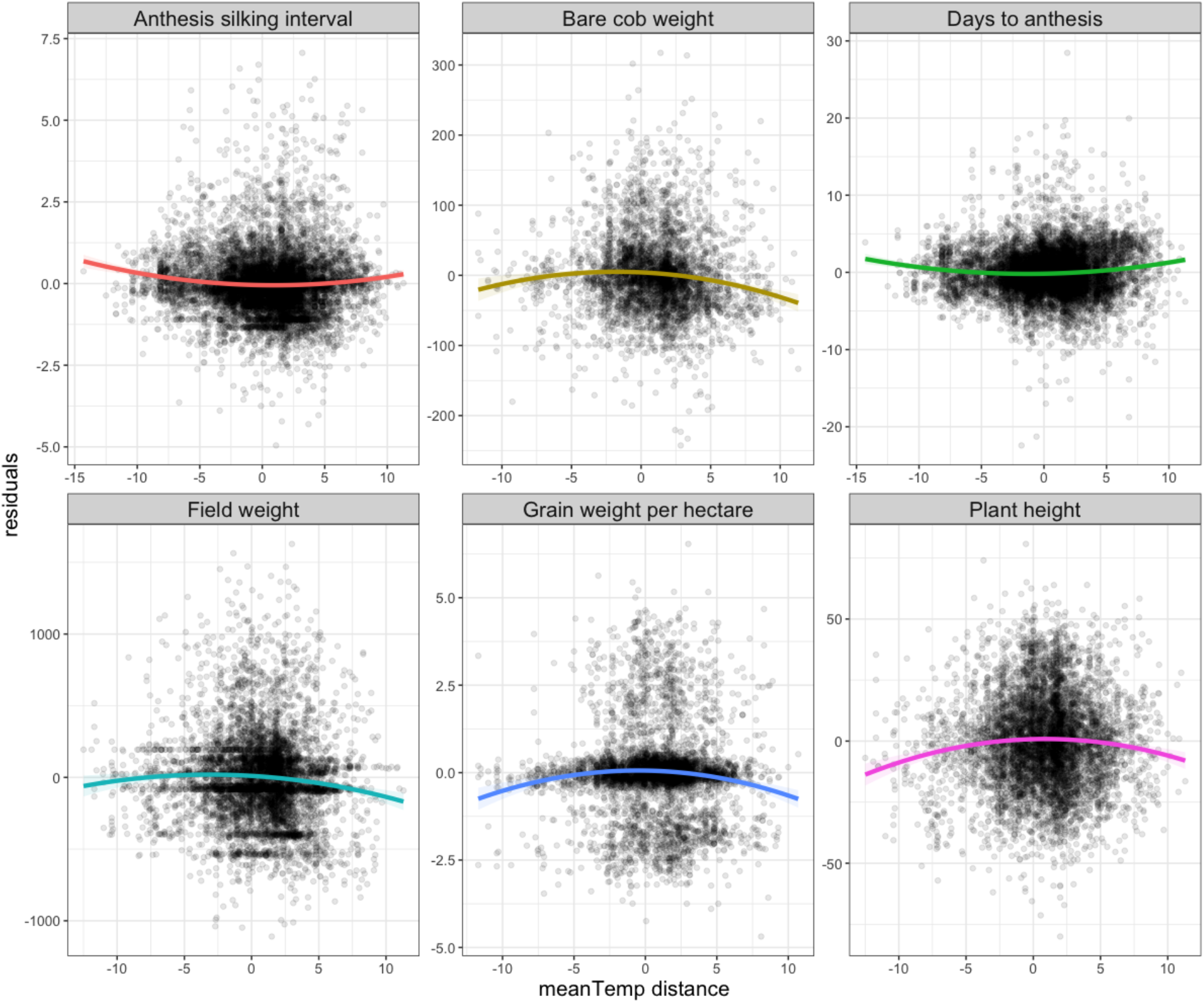
Plant phenotypes by difference in home and experimental trial temperature

**Figure S5.**
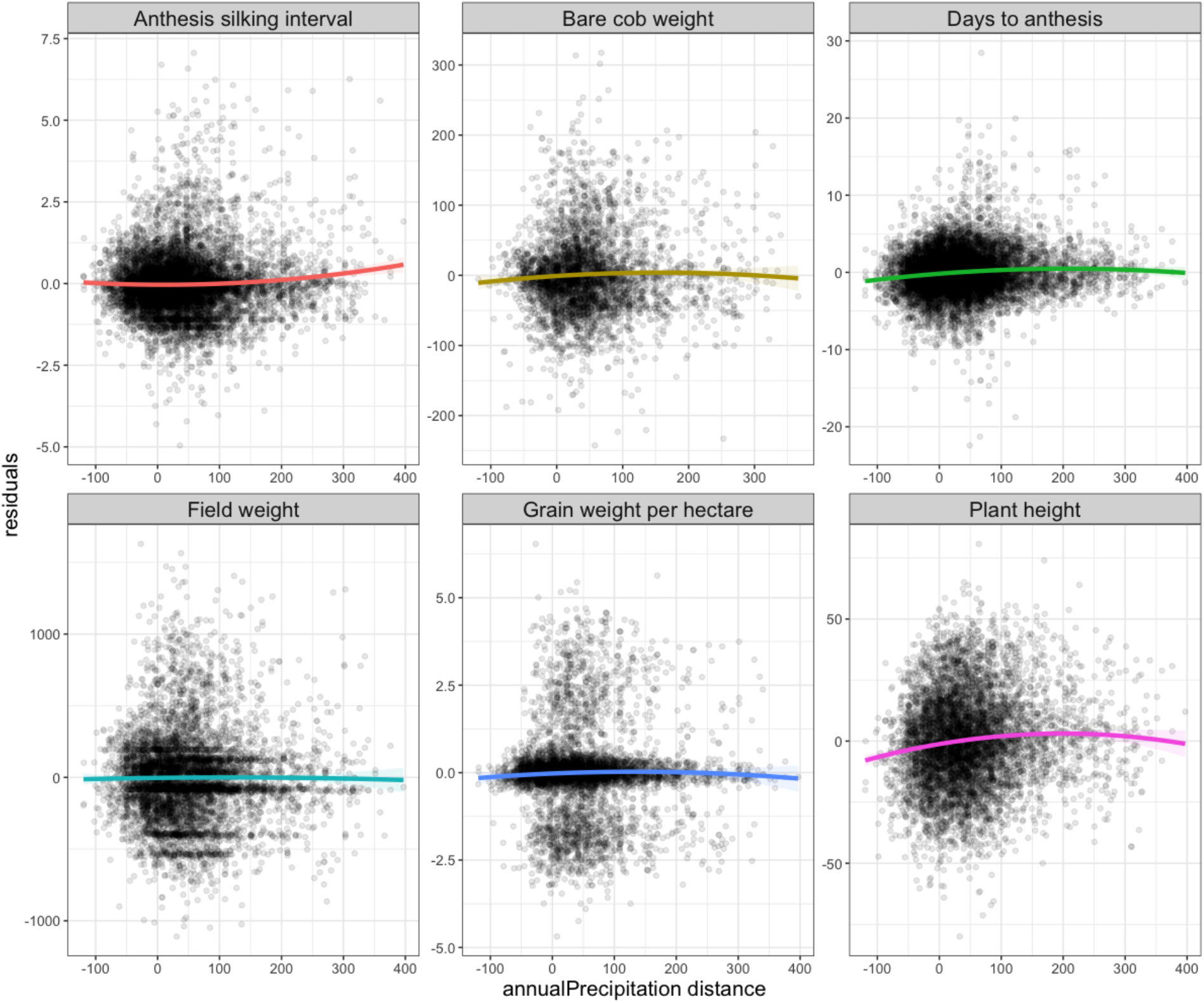
Plant phenotypes by difference in home and experimental trial precipitation

**Figure S6.**
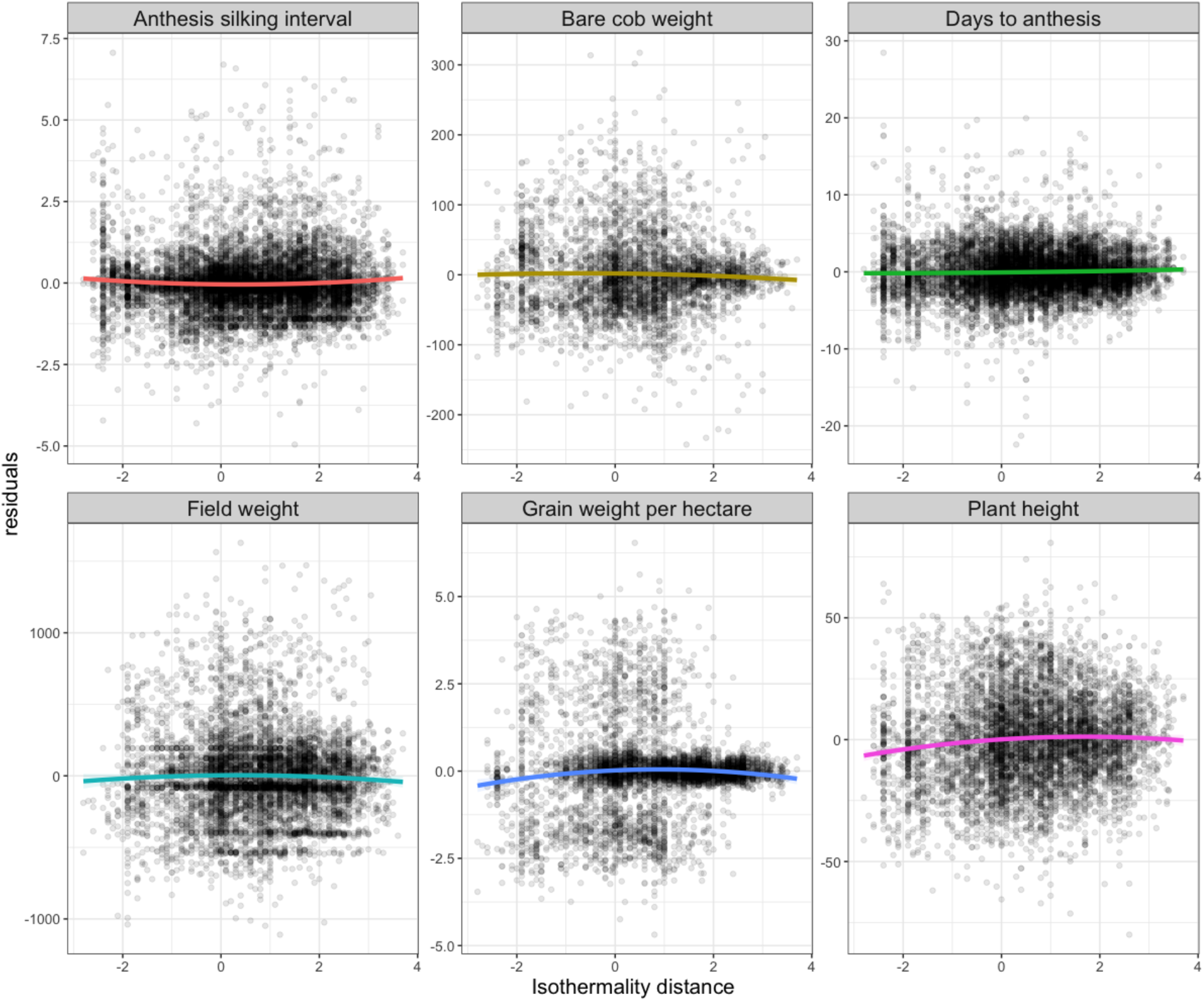
Plant phenotypes by difference in home and experimental trial isothermality

**Figure S7.**
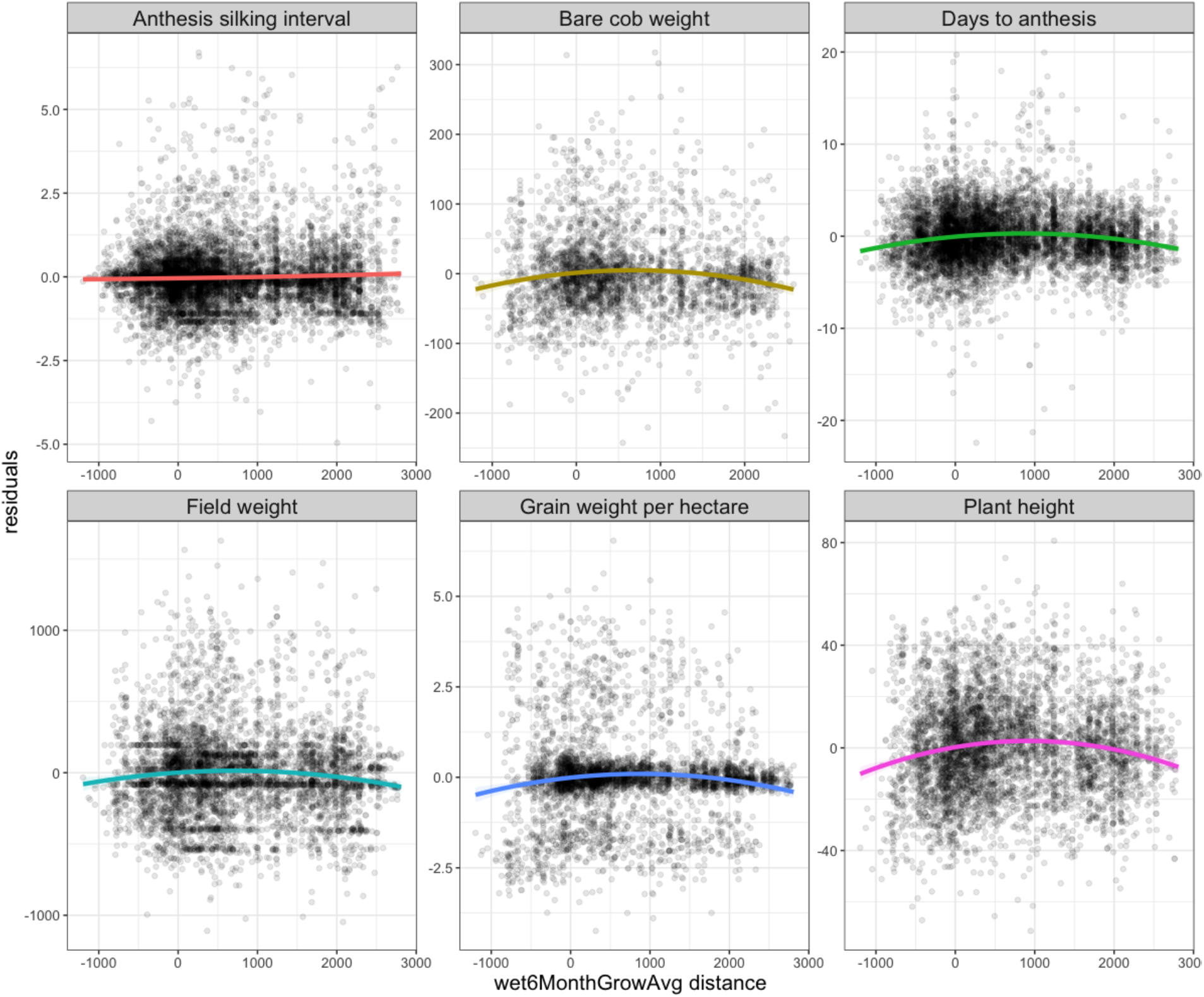
Plant phenotypes by difference in home and experimental trial wet days

**Figure S8.**
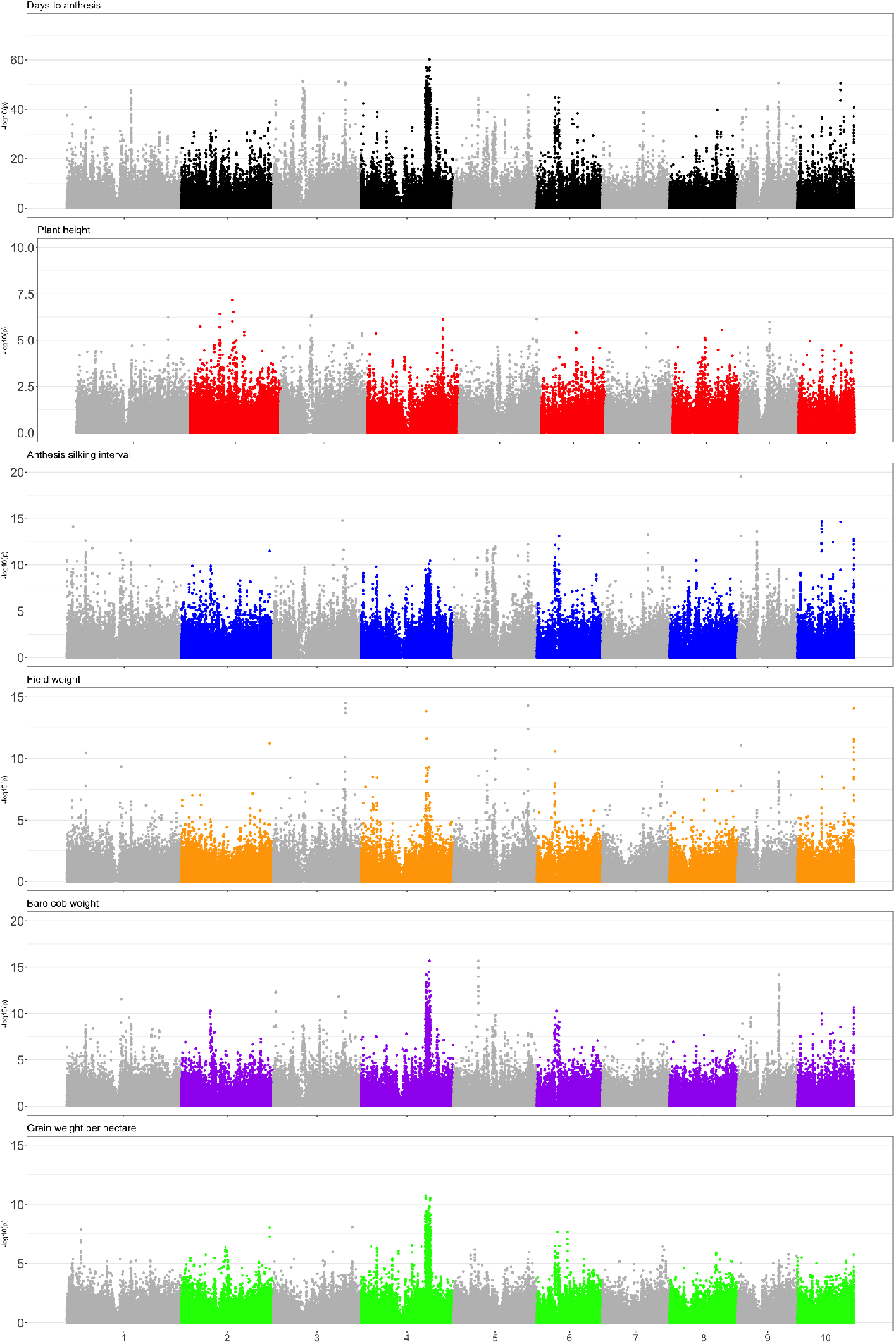
Manhattan plots for GxE GWAS analysis

**Figure S9.**
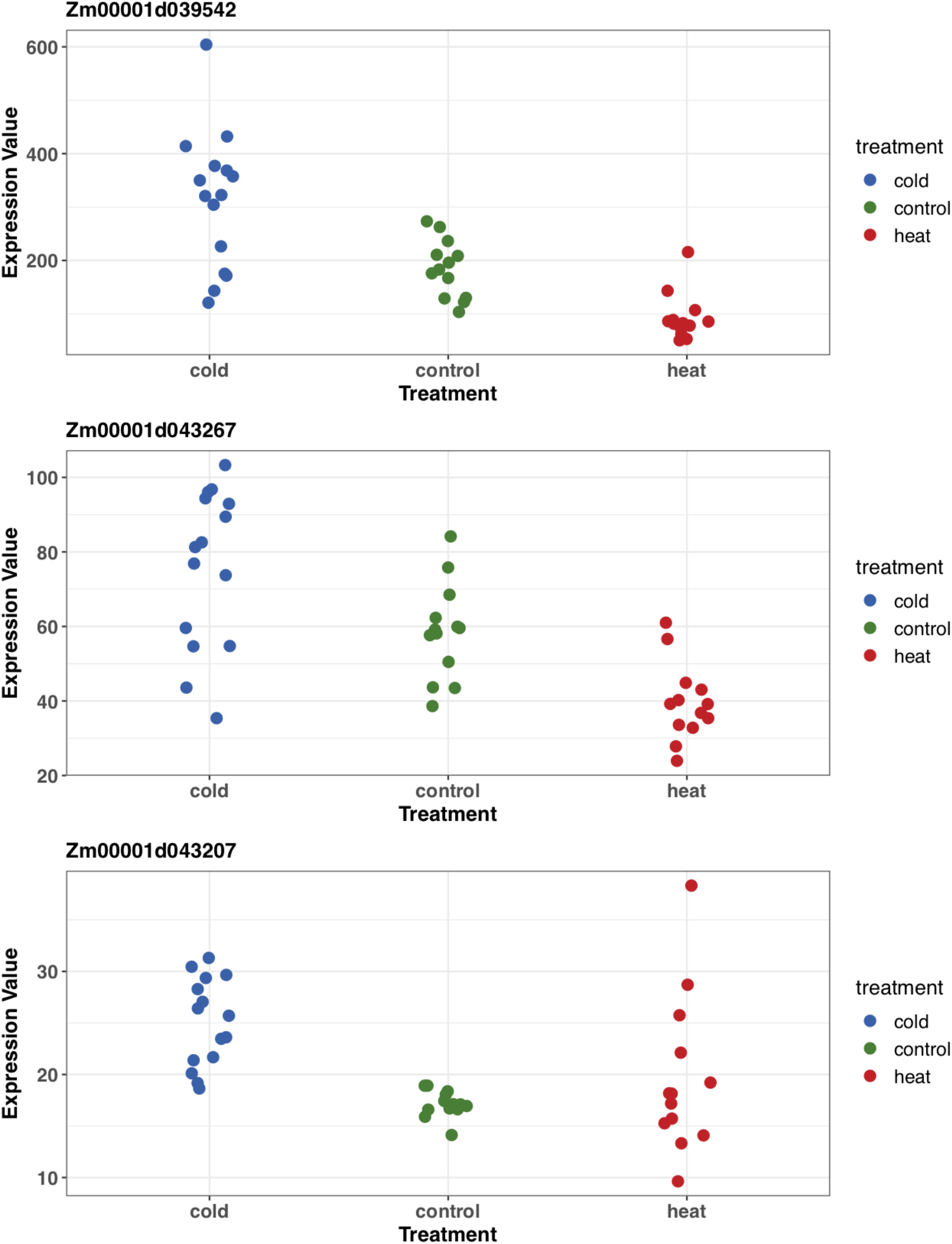
Expression values of lipid QTLxE candidate genes overlapping with GxE outliers. Data at each condition shows expression values of B73, Mo17, Oh43 and PH207 inbreds. Data was taken from (40).

**Figure S10.**
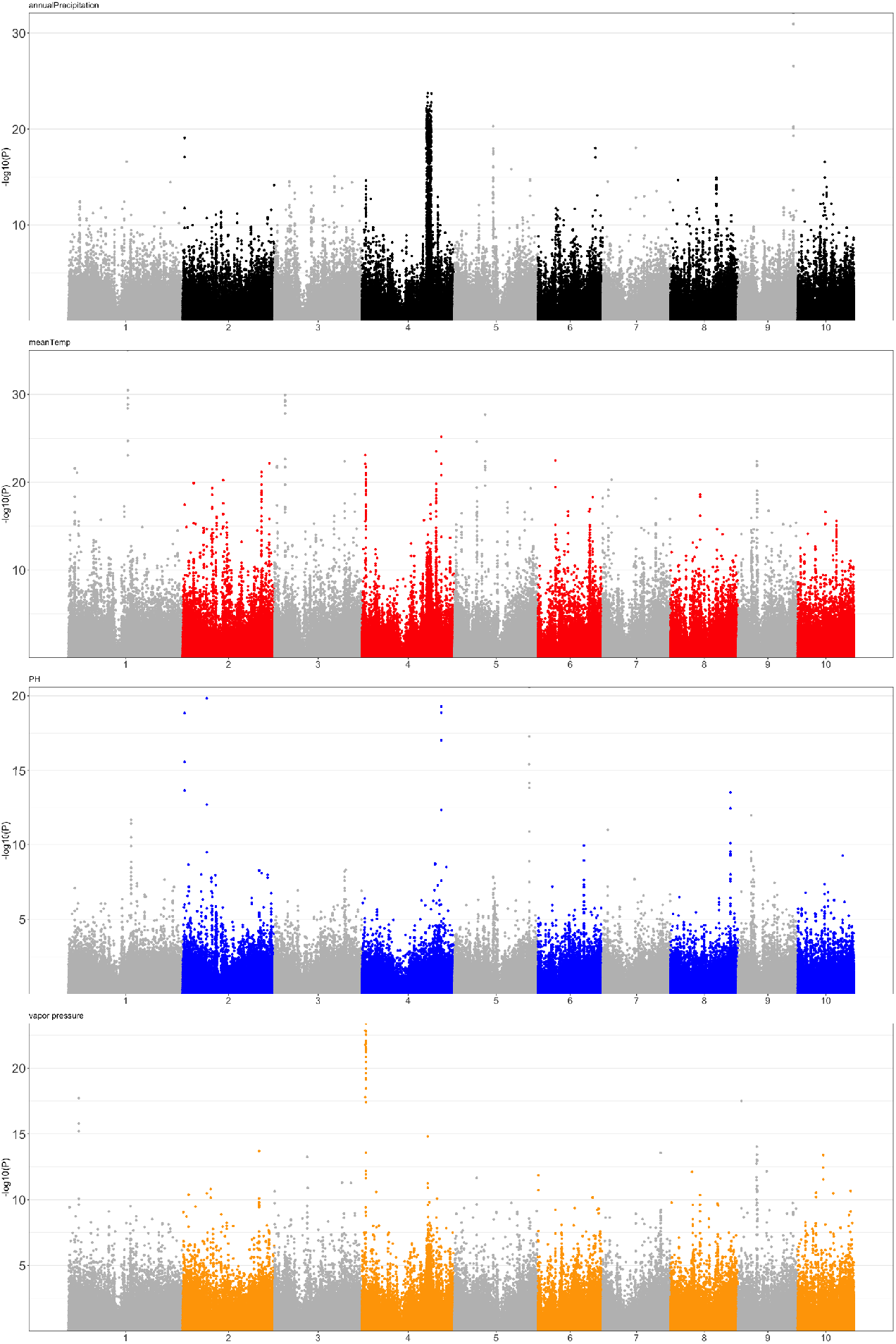
Manhattan plots for environmental GWAS analysis

**Figure S11.**
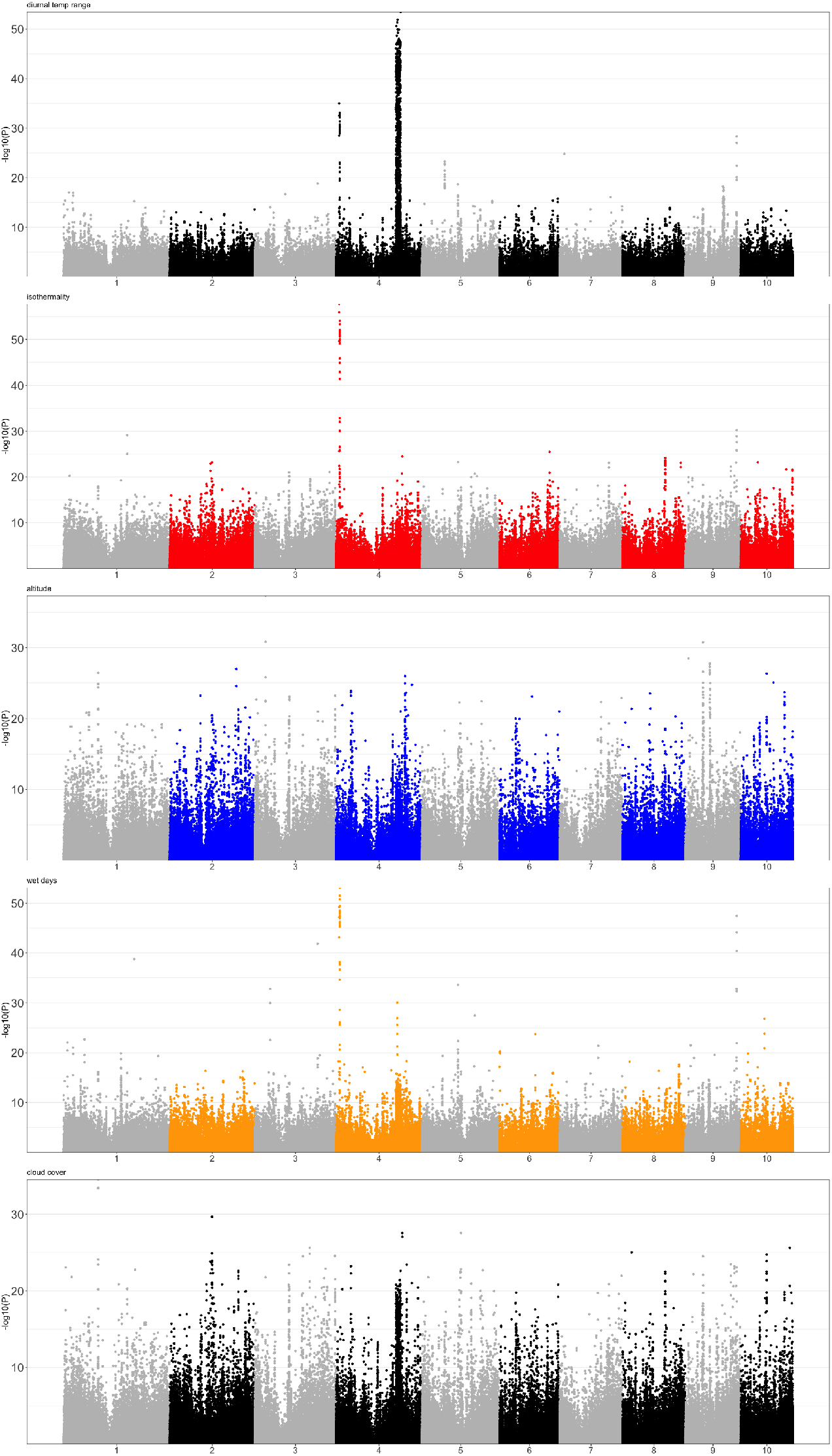
Manhattan plots for environmental GWAS analysis (continued)

**Figure S12.**
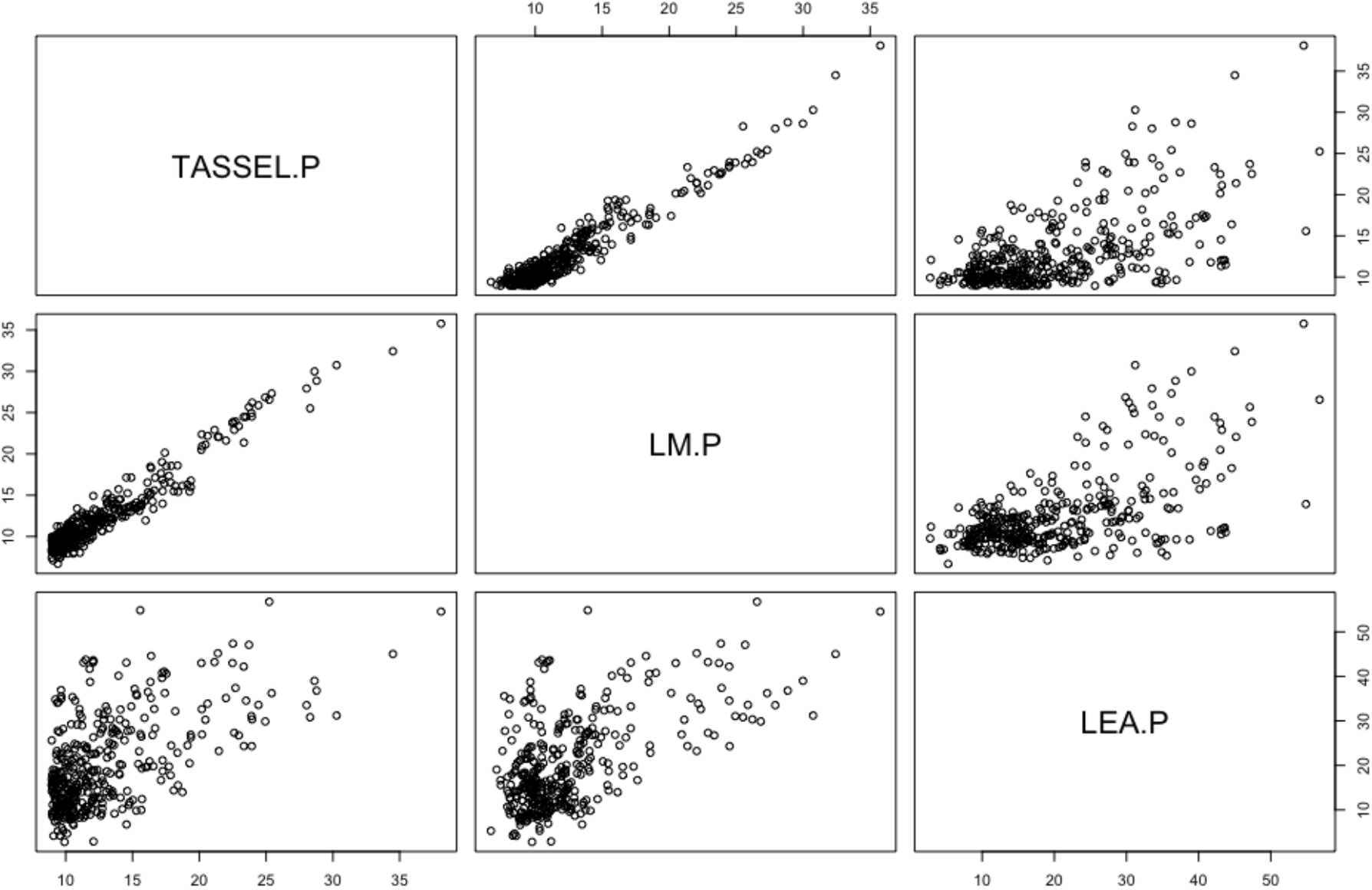
Pairwise comparisons of outliers calculated through different GWAS methods TASSEL, lm(R function), LFMM. All analyses use the first two PCs (TASSEL,lm) or two latent factors (LFMM) as a representative of population structure. TASSEL and lm are most similar likely reflecting differences in how latent factors and principal components account for population structure. All correlations are significant *P* < 2e^-16^

**Figure S13.**
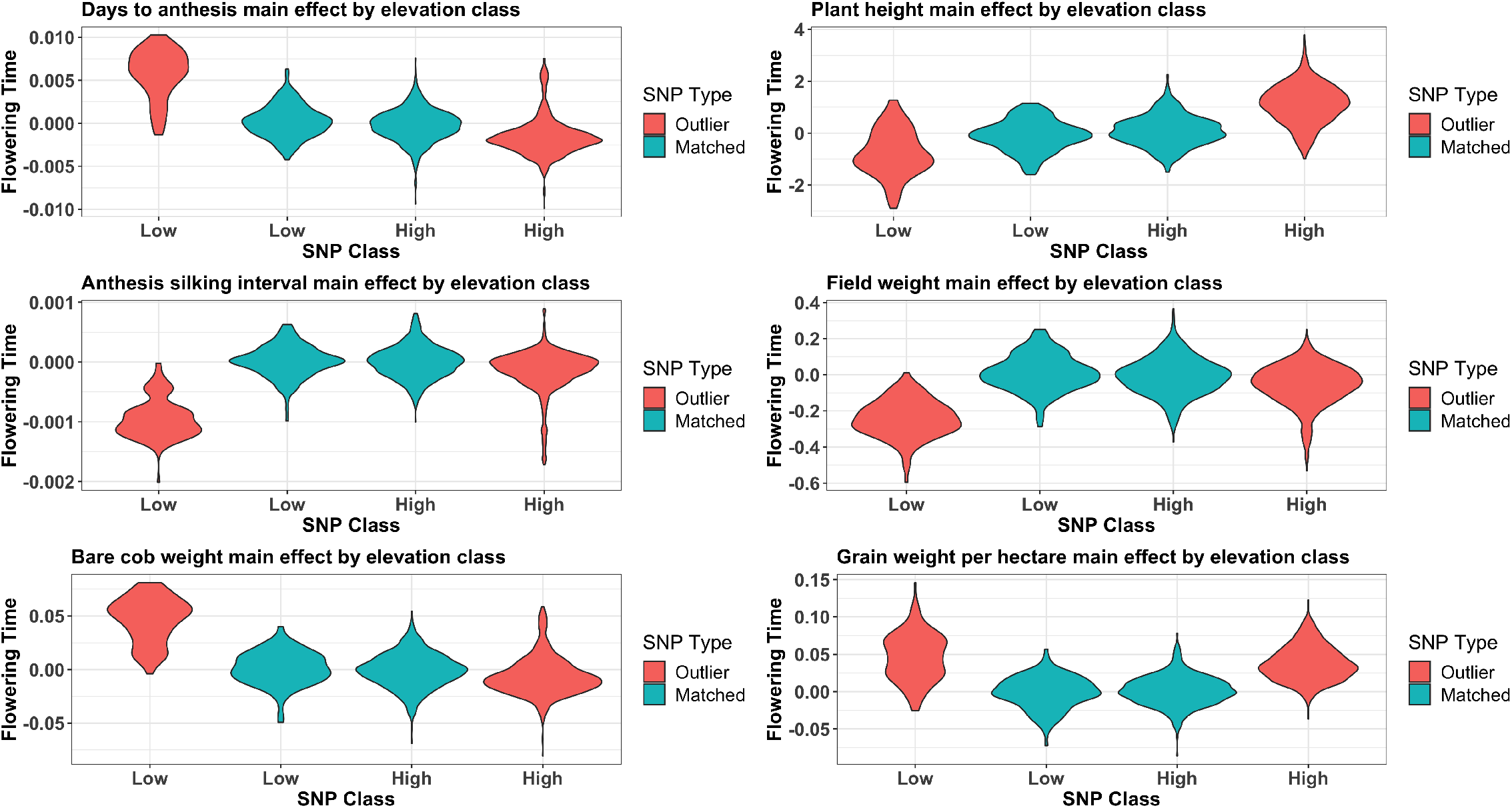
The main effect estimates of elevation associated alleles on phenotypes of the six different traits.

